# Allelopathic effect of *Anogeissus pendula* Edgew. and *Grewia flavescens* A.Juss on *Desmodium* species in Alwar district of Rajasthan

**DOI:** 10.1101/582023

**Authors:** Rajendra Prasad, A S Yadav

## Abstract

The effect of leaf leachates of *Anogeissus pendula* Edgew. and *Grewia flavescens* A.Juss was evaluated on growth of the three species of *Desmodium in* Alwar district of Rajasthan (27°4’ to 28°4’ N and 76°7’ to 77°13’ E). *Anogeissus pendula* reduced the seed germination of *Desmodium repandum* from 85% in control to 43% at 2% concentration, *Desmodium gangeticum* from 80% in control to 50% at 6% concentration and *Desmodium triflorum* from 78% in control to 28% at 0.1% concentration of leaf leachates. Similarly, the leaf leachates of *Grewia flavescens* reduced the seed germination of *Desmodium repandum* from 85% in control to 40% at 2% concentration, *Desmodium gangeticum* from 80% in control to 50% even at 0.1% concentration and *Desmodium triflorum* from 78% in control to 21% at 0.5% concentration. The leaf leachates of both the species also reduced the elongation of radicle and plumule of three Desmodium species; however, the adverse effect of leaf leachate of *Grewia flavescens* was more severe than that of the other species. Among the three *Desmodium* species, the allelopathic effect of *Grewia flavescens* and *Anogeissus pendula* was more severe on *Desmodium triflorum* as compared to the other two species. It may be suggested that the population of three species of *Desmodium* is partly regualated by the allelopathic effect of *Anogeissus pendula* and *Grewia flavescens* in this tropical dry deciduous forest.

## Introduction

Muller (1969) has suggested that competition as well as allelopathy comes in to play, such that, the pattern of herb distribution established at the time of germination is maintained throughout the growing season. The allelopathy phenomenon has received considerable attention as a fundamental ecological function (Rai and Tripathi 1984, Nakano *et al* 2003, 2004, Oudhia and Tripathi 1999, Ambika *et al*, 2003, Kohli et al 2004). Chung *et al* (2001) and Inderjit *et al* (2002) reported that allelochemicals and their mixture inhibited seed germination in grasses. Sharma *et al* (2009) investigated the effect of the extract of several weeds on seed germination of different varieties of wheat. *Desmodium repandum* (Vahl.) DC. is an annual herb which grows on the hill slopes, *Desmodium gangeticum (L.) DC.* a perennial robust herb grows commonly at the base of hill slopes and in the valleys and *Desmodium triflorum (L.) DC.* a perennial herb grows in disturbed forest areas and wastelands in Alwar district of Rajasthan (27°4’ to 28°4’ N and 76°7’ to 77°13’ E). The tropical dry deciduous thorn forest is dominated by a tree species, *Anogeissus pendula* and a shrub species, *Grewia flavescens* in this region. Both the species produce enormous amount of leaf litter which is added to the soil every year and it remains undecomposed until rains commences in July. The low population density (stems m^-2^) of *Desmodium repandum, Desmodium gangeticum* and *Desmodium triflorum* is 12, 4 and 9 respectively in their natural habitats (Prasad 2015) which indicates that associated vegetation may suppresses their growth. Hence, it may be inferred that these dominant species may adversely affect the growth of *Desmodium* species through leachates released from leaf litter in the rainy season. An attempt has, therefore, been made in the present study to evaluate the allelopathic effect of *Anogeissus pendula* and *Grewia flavescens* on seed germination and seedling growth of the three species of *Desmodium* which commonly grow in Alwar district of Rajasthan.

## Materials and Methods

Fallen dry leaves of *Anogeissus pendula* and *Grewia flavescens* were collected from their natural growing sites. The aqueous leaf leachates of 0.1, 0.2, 0.5, 1.0, 2.0, 4.0, 6.0 percent concentration were prepared. 100 mg, 200 mg, 500 mg, 1.0 gm, 2.0 gm, 4.0 gm and 6.0 gm leaves of *Anogeissus pendula* were soaked for 48 hours in 50 ml distilled water in separate beakers. 100 mg, 200 mg, 500 mg, 1.0 gm, 2.0 gm, 4.0 gm and 6.0 gm leaves of *Grewia flavescens* were also soaked for 48 hours in 50 ml distilled water in other separate beakers. Aqueous solution of leachates was filtered through a muslin cloth separately for both the species and each concentration. Each of these aqueous leachates was made to 100 ml by adding distilled water. Before setting seed germination experiment, *Desmodium gangeticum* and *Desmodium triflorum* seeds were treated with H_2_SO_4_ for one minute to break the seed coat dormancy and then thoroughly washed with distilled water. The effect of leaf leachates on seed germination was studied by soaking the seeds of *Desmodium repandum, Desmodium gangeticum* and *Desmodium triflorum* in different concentrations for 24 hours. Thirty soaked seeds were kept for germination on filter paper underlined by cotton moistened with the respective leaf leachate solution in the three petridishes for each concentration of every dominant plant species. The petridishes were kept for seed germination at ± 30° C temperature in a seed germinator and observations were taken. A control was run using distilled water as soaking medium.

Aqueous leachates of leaves of *Anogeissus pendula* and *Grewia flavescens* in distilled water were also tested for their effect on radical and plumule growth of *Desmodium repandum*, *Desmodium gangeticum* and *Desmodium triflorum*. Leachates solution of 0.1, 0.2, 0.5, 1.0, 2.0, 4.0, 6.0 percent concentration were used. For studying the effect of leachates on growth of *Desmodium* species length of seedlings radicle and plumule were measured after 15 days with the help of a calliper.

## Results and Discussion

The leaf leachates of *Anogeissus pendula* reduced the seed germination of *Desmodium repandum* from 85% in control to 43% at 2% concentration of leaf leachates (Table 1). The adverse effect of leaf leachates of *Anogeissus pendula* on seed germination and seeding growth was from 0.5% concentration of onwards. The leaf leachates of *Anogeissus pendula* have less effect on seedling growth as the reduction in radicle elongation was from 6.4 cm in control to 4.1 cm at 2% concentration while the reduction in plumule elongation was slightly less than radicle elongation. *Grewia flavescens* reduced seed germination of *Desmodium repandum* from 85% in control to 40% at 2% concentration while the elongation of radicle from 6.4 cm to 4.8 cm and plumule elongation from 3.4 cm to 2.24 cm with increase in concentration of leaf leachates (Table 1). This is in conformity with Dhawan and Dhawan (1995) who reported the adverse effect of leaf extract of *Cassia tora* on *Parthenium hysterophorus*. Gupta and Yadav (2007) also reported that leaf leachates of *Cassia tora* inhibits seed germination and seedling growth of the grass species of the Sariska Tiger Reserve forest. Similar results were obtained by Dey and Yadav (2010) who evaluated the allelopathic effect of exotic and indigenous species on seed germination and seedling growth of *Indigofera trita*. However, Dey and Yadav (2010) observed that the leaf leachates of *Holoptelia integrifolia,* a local tree of this region, have little effect on seed germination of *Indigofera trita*. The harmful effect on radicle and plumule elonagation begins after 0.2% concentration of leaf leachates in this species. However, the intensity of adverse effect of *Grewia flavescens* was slightly higher on seed germination and seedling growth of *Desmodium repandum* than that of leaf leachates of *Anogeissus pendula*.

**Table 1.** Allelopathic effect of leaf leachates of Anogeissus pendula and Grewia flavescens on seed germination and seedling growth of Desmodium repandum (±SE).

| Leaf leachates | Concentration of Leaf leachates (%) | Seed germination (%) | Radicle elongation(cm) | Plumule elongation(cm) |
| --- | --- | --- | --- | --- |
| *Anogeissus pendula* | 0.1 | 74.99±5.90 | 5.64±0.14 | 3.34±0.17 |
| | 0.2 | 63.33±11.79 | 5.5±0.2 | 3.2±0.23 |
| | 0.5 | 48.33±10.61 | 5.14±0.09 | 2.46±0.15 |
| | 1 | 46.67±9.43 | 4.44±0.15 | 2.34±0.17 |
| | 2 | 43.33±7.07 | 4.14±0.09 | 2.2±0.12 |
| *Grewia flavescens* | 0.1 | 66.66±4.71 | 5.9±0.20 | 2.9±0.48 |
| | 0.2 | 61.66±10.61 | 5.5±0.16 | 2.84±0.24 |
| | 0.5 | 56.83±4.83 | 5.44±0.28 | 2.7±0.23 |
| | 1 | 48.33±1.18 | 5.24±0.22 | 2.6±0.26 |
| | 2 | 40±7.07 | 4.84±0.24 | 2.24±0.10 |
| Control | Distilled Water | 84.99±1.18 | 6.4±0.52 | 3.4±0.38 |

Incase of *Desmodium gangeticum,* the adverse effect of leaf leachates of *Anogeissus pendula* was higher on seed germination than on seedling growth (Table 2). The seed germination decreased from 80% in control to 50% at 6% concentration of leaf leachates. Similarly the radicle and plumule elogation is reduced from 3.1 to 2.0 cm and 1.8 cm to 0.7 cm, respectively with increase in concentration. The growth of seedlings was adversely affected only at higher concentration of leaf leachates of *Anogeissus pendula,* however the reduction of plumule elongation was more as compared to radicle elongation. This is in agreement with Dey and Yadav (2010) who also reported that the adverse effect allelochemics was more on plumule than on radicle elongation in *Indigofera trita*. The leaf leachates of *Grewia flavescens* caused greater reduction in seed germination of *Desmodium gangeticum* from 80% in control to 50% even at 0.1% concentration which was further reduced to 10% at 6% concentration of leachates (Table 2). Similarly the radicle and plumule elongation was also reduced from 3.1 cm to 1.5 cm and 1.8 cm to 0.7 cm, respectively with the increase in concentration of leaf leachates up to 6%. In this case also, the plumule elogation is more sensitive to leaf leachates than the radicle elongation.

**Table 2.** Allelopathic effect of leaf leachates of Anogeissus pendula and Grewia flavescens seed germination and seedling growth of Desmodium gangeticum (±SE).

| Leaf leachates | Concentration of leaf leachates (%) | Seed germination (%) | Radicle elongation (cm) | Plumule elongation (cm) |
| --- | --- | --- | --- | --- |
| *Anogeissus pendula* | 0.1 | 61.66±3.54 | 3±0.14 | 1.6±0.15 |
| | 0.2 | 59.99±4.71 | 2.86±0.09 | 1.36±0.12 |
| | 0.5 | 63.33±2.35 | 2.66±0.17 | 1.24±0.12 |
| | 1 | 65.16±1.06 | 2.64±0.17 | 1.08±0.27 |
| | 2 | 56.66±2.36 | 2.24±0.10 | 0.96±0.23 |
| | 4 | 53.33±4.72 | 2.14±0.10 | 0.84±0.18 |
| | 6 | 49.99±2.36 | 1.98±0.14 | 0.74±0.19 |
| *Grewia flavescens* | 0.1 | 49.99±2.36 | 2.9±0.17 | 1.38±0.19 |
| | 0.2 | 46.66±2.36 | 2.68±0.10 | 1.32±0.20 |
| | 0.5 | 28.33±1.18 | 2.54±0.12 | 1.24±0.17 |
| | 1 | 24.99±1.18 | 2.42±0.15 | 1.08±0.14 |
| | 2 | 23.33±2.35 | 2.2±0.17 | 0.92±0.19 |
| | 4 | 18.33±1.18 | 1.9±0.17 | 0.82±0.13 |
| | 6 | 9.99±2.36 | 1.52±0.16 | 0.68±0.10 |
| Control | Distilled water | 79.99±2.36 | 3.1±0.17 | 1.76±0.22 |

Leaf leachates of *Anogeissus pendula* reduced seed germination of *Desmodium triflorum* from 78% in control to 28% at 0.1% concentration which was further reduced to complete inhibition of germination with increase in leaf leachates concentration from 0.1% to 6.0% (Table 3). Similarly, the elongation of radicle was reduced from 2.8 cm to 2.2 cm at 0.1% concentration which further decreased to nil with increase in 0.2% to 6.0% concentration of leaf leachates. The adverse effect of leaf leachates of *Grewia flavescens* was more severe on the seed germination and seedling growth of *Desmodium triflorum* (Table 3). The seed germination was reduced from 78% in control to 21% at 0.5% concentration and at more than 2% there was complete inhibition of seed germination. Similar adverse effect of leaf leachates was observed on elongation of radicle and plumule of this species. These observations suggest that the leaf leachates of the shrub, *Grewia flavescens,* have more adverse effect on seed germination and seedling growth of the three species of *Desmodium.* Among the three *Desmodium* species, the allelopathic effect of *Grewia flavescens* and *Anogeissus pendula* was more severe on the seed germination and seedling growth of *Desmodium triflorum.* Dey and Yadav (2010) have also observed that the leaf leachates of indigenous shrub, *Capparis sepiaria* exhibited higher inhibitory effect on seed germination and seedling growth of *Indigofera trita* than that of *Holoptelia integrifolia* a tree species of this region. It may be concluded that the growth and distribution of three species of *Desmodium* is partly regulated by the allelopathic effect of *Anogeissus pendula* and *Grewia flavescens* in the tropical dry deciduous forest of Alwar district in Rajasthan.

**Table 3.** Allelopathic effect of leaf leachates of Anogeissus pendula and Grewia flavescens seed germination and seedling growth of Desmodium triflorum (±SE).

| Leaf leachates | Concentration of Leaf leachates (%) | Seed germination (%) | Radicle elongation(cm) | Plumule elongation(cm) |
| --- | --- | --- | --- | --- |
| *Anogeissus pendula* | 0.1 | 28.33±1.18 | 2.26±0.14 | 1.5±0.16 |
| | 0.2 | 18.33±1.18 | 1.8±0.23 | 1.34±0.15 |
| | 0.5 | 16.66±2.36 | 1.44±0.19 | 1.16±0.13 |
| | 1 | 14.99±1.18 | 1.34±0.26 | 1±0.15 |
| | 2 | 11.67±1.18 | 1.14±0.25 | 0.82±0.11 |
| | 4 | 8.33±1.18 | 0.82±0.19 | 0.74±0.1 |
| | 6 | 0±0 | 0±0 | 0±0 |
| *Grewia flavescens* | 0.1 | 46.665±2.36 | 2.12±0.16 | 1±0.15 |
| | 0.2 | 34.995±1.18 | 1.88±0.21 | 0.94±0.18 |
| | 0.5 | 21.665±1.18 | 1.78±0.26 | 0.9±0.18 |
| | 1 | 14.995±1.18 | 1.68±0.19 | 0.76±0.20 |
| | 2 | 9.995±2.36 | 1.14±0.29 | 0.66±0.17 |
| | 4 | 0±0 | 0±0 | 0±0 |
| | 6 | 0±0 | 0±0 | 0±0 |
| Control | Distilled water | 78.33±1.18 | 2.76±0.22 | 1.56±0.28 |

## Acknowledgements

The financial assistance received from University grants Commission, New Delhi as a Minor Research Project by the first author is gratefully acknowledged.

